# OptoGranules reveal the evolution of stress granules to ALS-FTD pathology

**DOI:** 10.1101/348870

**Authors:** Peipei Zhang, Baochang Fan, Peiguo Yang, Jamshid Temirov, James Messing, Hong Joo Kim, J. Paul Taylor

## Abstract

Stress granules are non-membranous assemblies of mRNA and protein that form in response to a variety of stressors. Genetic, pathologic, biophysical and cell biological studies have implicated disturbances in the dynamics of membrane-less organelles, such as stress granules, as a pathobiological component of amyotrophic lateral sclerosis (ALS) and frontotemporal dementia (FTD)^1–12^. This confluence of evidence has inspired the hypothesis that these diseases reflect an underlying disturbance in the dynamics and material properties of stress granules; however, this concept has remained largely untestable in available models of stress granule assembly, which require the confounding variable of exogenous stressors. Here we demonstrate the development and use of a light-inducible stress granule system, termed OptoGranules, which permits discrete, experimental control of the dynamics and material properties of stress granules in living cells in the absence of exogenous stressors. The nucleator in this system is Opto-G3BP1, a light-sensitive chimeric protein assembled from the intrinsically disordered region (IDR) and RNA-binding domain of G3BP1 combined with the light-sensitive oligomerization domain of *Arabidopsis thaliana* cryptochrome 2 (CRY2) photolyase homology region (PHR). Upon stimulation with blue light, Opto-G3BP1 initiates the rapid assembly of dynamic, cytoplasmic, liquid granules that are composed of canonical stress granule components, including G3BP1, PABP, TIA1, TIAR, eIF4G, eIF3η, ataxin 2, GLE1, TDP-43 and polyadenylated RNA. With this system, we demonstrate that persistent or repetitive assembly of stress granules is cytotoxic and is accompanied by the evolution of stress granules to neuronal cytoplasmic inclusions that recapitulate the pathology of ALS-FTD.

---

Membrane-less organelles such as stress granules and related RNA granules are microscopically visible, mesoscale structures that arise through liquid-liquid phase separation (LLPS), a biophysical phenomenon in which RNA-protein complexes separate from the surrounding aqueous cytoplasm to create a functional cellular compartment with liquid properties. The assembly of RNA granules is driven by the collective behavior of many types of macromolecular interactions, including RNA-RNA interactions, protein-RNA interactions, conventional interactions between folded protein domains, as well as weak, transient interactions mediated by low complexity, intrinsically disordered regions (IDRs) of proteins - particularly those present in RNA-binding proteins^13^.

G3BP1 (and its close paralog G3BP2) is an essential nucleator of stress granule assembly^14^. G3BP1 has an N-terminal 142-amino acid dimerization domain, termed the NTF2 domain, that is essential for nucleation of stress granule assembly. Remarkably, the NTF2 domain can be replaced by generic dimerization domains, and the resulting chimeric proteins are able to fully nucleate stress granule assembly in living cells (Yang, in preparation). Thus, the domain architecture of G3BP1 is ideal for engineering light-inducible stress granule assembly by replacing the NTF2 domain of G3BP1 with the blue light-dependent dimerization domain CRY2_PHR_ in frame with the fluorescent protein mCherry. We named this construct “Opto-G3BP1,” and also created an “Opto-Control” construct referring to CRY2_PHR_-mCherry alone (**Fig. 1a**). Consistent with prior reports, knockout of endogenous *G3BP1* and *G3BP2* in U2OS cells abolished stress granule assembly in response to arsenite^14^ (**Extended Data Fig. 1a**). When introduced to these *G3BP1/G3BP2* double knockout cells, Opto-G3BP1 (as well as the analogous chimeric protein Opto-G3BP2) significantly restored stress granule assembly in response to arsenite, demonstrating that these chimeric proteins are functionally intact (**Extended Data Fig. 1b-e**).

**Figure 1.**
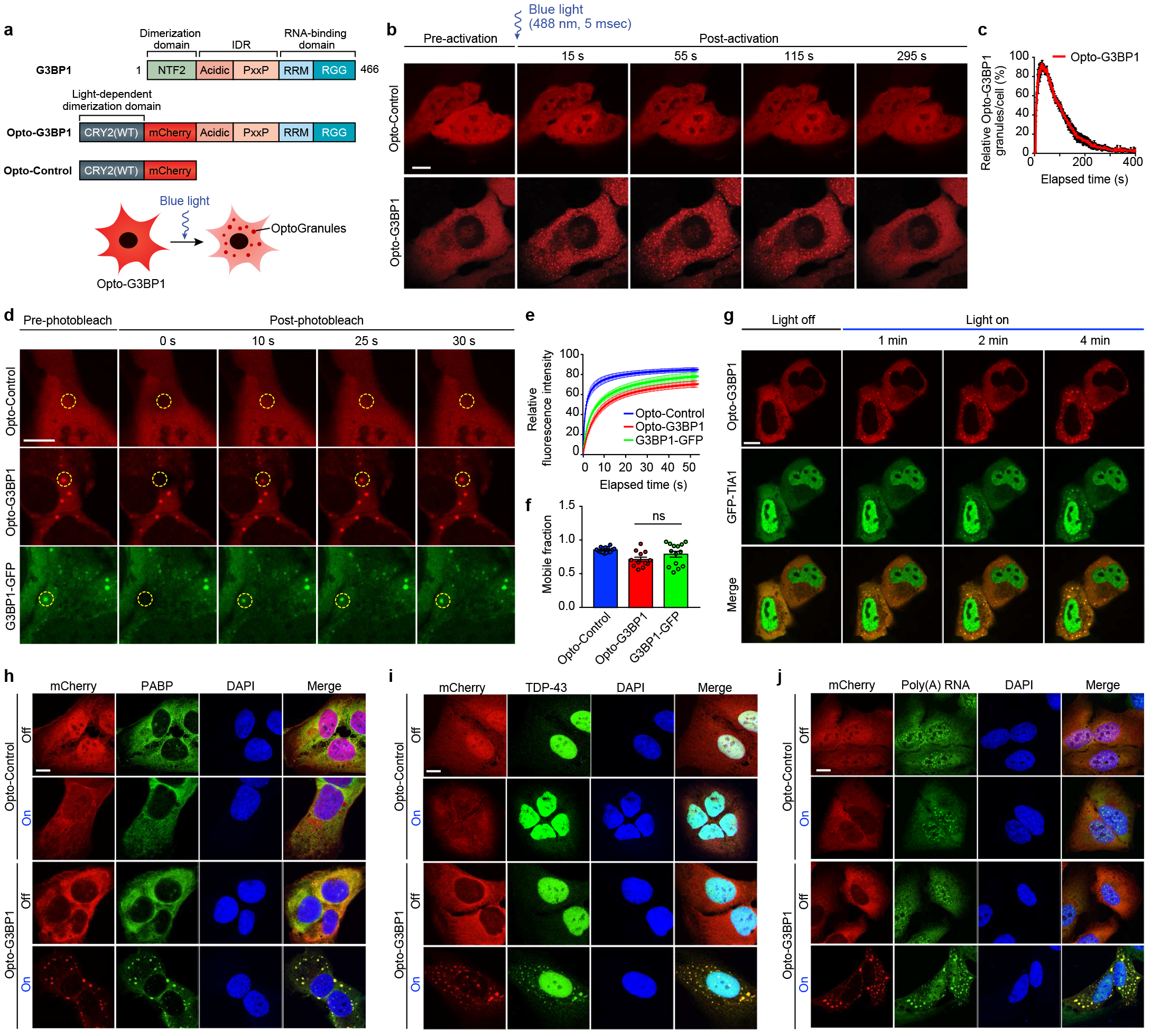
OptoGranules are light-inducible dynamic stress granules. **a**, Design of Opto-G3BP1 and Opto-Control constructs. **b,** U2OS cells stably expressing Opto-Control or Opto-G3BP1 were stimulated with a single 5-msec pulse of 488-nm blue light. Representative images are shown from n = 3 independent experiments. **c,** Quantification of data in cells treated as in **b**. Five cells with similar expression levels were counted. Granule numbers are shown relative to the granule number at the peak of OptoGranule assembly. Error bars represent s.e.m. **d-f,** U2OS cells were stably transfected with Opto-Control or Opto-G3BP1, or stable Opto-G3BP1 cells were transiently transfected with G3BP1-GFP, and stimulated with blue light for 3 mins. Regions marked with yellow circles were photobleached and monitored for fluorescence recovery. Data are shown as representative images (**d**), relative fluorescence intensity of photobleached region over time (**e**), and relative mobile fraction derived from **e** (**f**). For **e,f**, n = 15 cells for Opto-Control; n = 12 for Opto-G3BP1; n = 14 for G3BP1-GFP. Data are representative of n = 3 independent experiments. Data shown as mean + s.e.m. ns, not significant by one-way ANOVA with Dunnett’s post test. **g,** U2OS cells were transiently transfected with Opto-G3BP1 and the stress granule marker GFP-TIA1 and stimulated with blue light. Representative images are shown from n = 3 independent experiments. **h-j,** U2OS cells stably expressing Opto-Control or Opto-G3BP1 constructs without or with blue light stimulation were immunostained with PABP antibody (**h**), TDP-43 antibody (**i**), or RNA fluorescence in situ hybridization using FAM-labelled oligo (dT)20 as a probe (**j**). Scale bars, 10 μm in all micrographs.

We next generated U2OS cell lines stably expressing Opto-G3BP1 or Opto-Control constructs. Within seconds of blue light activation, Opto-G3BP1 in U2OS cells assembled into cytoplasmic granules, which we termed OptoGranules (**Fig. 1b and Supplementary Videos 1, 2**).

Remarkably, a 5-millisecond pulse of 488-nm blue light was sufficient to initiate robust induction of cytoplasmic granules, and these granules spontaneously disassembled over a period of approximately 5 minutes (**Fig. 1b,c**). These granules are highly dynamic, exhibiting liquid behaviors such as fusion to form larger granules and relaxation to a spherical shape (**Supplementary Video 2**). In contrast, under the same conditions, Opto-Control expression remained diffuse, with a modest amount of nuclear and cytoplasmic clusters (**Fig. 1b and Supplementary Video 1**). To confirm the reversible nature of OptoGranules, we performed fluorescence recovery after photobleaching (FRAP) to monitor recovery rates and mobile fractions of individual granules (**Fig. 1d-f**), finding that these properties were very similar between Opto-G3BP1 and the conventional stress granules marker G3BP1-GFP.

To further define the relationship of OptoGranules to stress-induced stress granules, we next examined the composition of OptoGranules. Employing live cell imaging, we documented the dynamic recruitment of the stress granule marker GFP-TIA1 into OptoGranules following light-induced assembly (**Fig. 1g and Supplementary Video 3**). Moreover, all stress granule components that we examined, including G3BP1, PABP, TDP-43, TIA1, TIAR, eIF4G, eIF3h, ataxin 2 and GLE1 were recruited to OptoGranules (**Fig. 1h-i** and **Extended Data Fig. 1f-k**). Since stress granules represent assembly of mRNA as well as protein^15,16^, we used fluorescent *in situ* hybridization (FISH) with fluorescently conjugated oligo(dT) probes to examine whether polyadenylated mRNAs are present in OptoGranules as in canonical stress granules. We found that polyadenylated mRNAs were recruited into OptoGranules that assembled after blue light stimulation but showed no relocalization in cells expressing Opto-Control (**Fig. 1j**). These findings indicate that OptoGranules are stress granules composed of mRNAs and RNA-binding proteins, including ALS-associated proteins such as TDP-43, ataxin 2, GLE1 and TIA1.

These results contrast with observations made with light-induced aggregation of so-called “OptoDroplets” that nicely illustrate intracellular phase transitions of proteins harboring IDRs, but do not represent assembly of complex, physiologically relevant membrane-less organelles^17^. Indeed, we directly examined the difference between OptoDroplets and OptoGranules. When expressed in U2OS cells, Opto-FUS [CRY2_PHR_-mCherry-FUS(IDR)] and Opto-TDP-43 [CRY2_PHR_-mCherry-TDP-43(IDR)] did assemble into droplets with blue light activation, as previously reported^17^, but these OptoDroplets did not colocalize with the stress granule marker G3BP1-GFP (**Extended Data Fig. 2a-b,d-e**). Similarly, constructs containing the IDR and RNA recognition motifs of TDP-43 or FUS [CRY2_PHR_-mCherry-FUS (1-371aa); CRY2_PHR_-mCherry-TDP-43 (106-414aa)] assembled into droplets upon blue light activation, but these droplets were negative for the stress granule marker PABP (**Extended Data Fig. 2c,f**). Expression of Opto-constructs using full-length TDP-43 or FUS [CRY2_PHR_-mCherry-TDP-43(FL); CRY2_PHR_-mCherry-TDP-43(FL)] did not produce cytoplasmic clusters with blue light activation (**Extended Data Fig. 2c,f**). Finally, Opto-TIA1, which represents fusion of CRY2 with the well-known stress granule protein TIA1 (CRY2_PHR_-mCherry-TIA1), also assembled into droplets with blue light activation, but did not drive the assembly of stress granules, as illustrated by lack of colocalization with G3BP1-GFP (**Extended Data Fig. 2g**). These data indicate that the formation of FUS-, TDP-43-, or TIA1-based OptoDroplets does not recapitulate the formation of stress granules, and is consistent with the proposition that a specific nucleator is required to initiate the assembly of distinct membrane-less organelles^13^.

**Figure 2.**
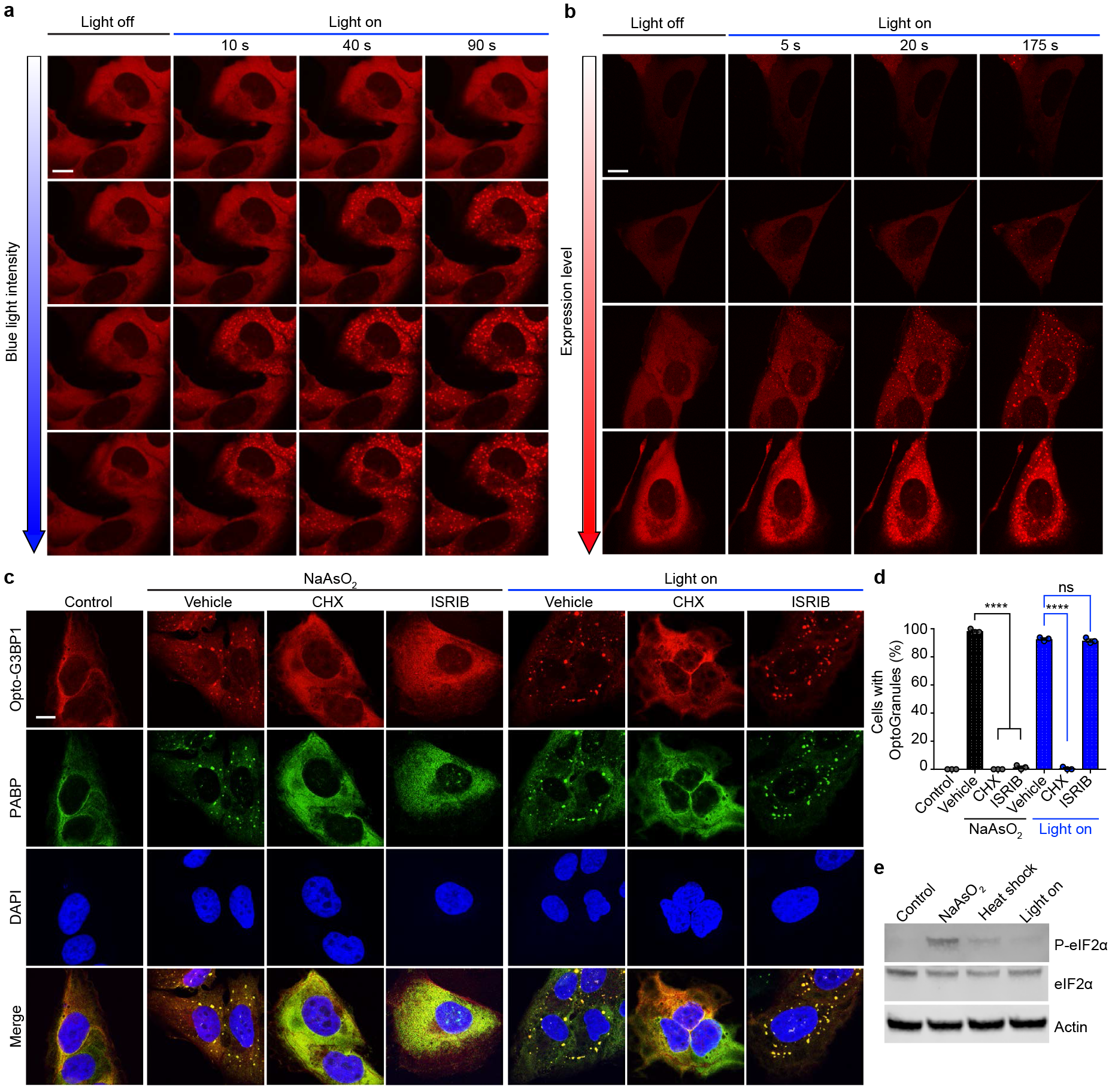
OptoGranule formation is dependent on the local concentration of activated G3BP1 and dependent on polysome disassembly, but independent of eIF2α phosphorylation. **a,** U2OS cells stably expressing Opto-G3BP1 were exposed to blue light of increasing intensity (488 nm laser power and energy measurement from top to bottom W/cm^2^: 1%, 0.0243; 5%, 0.0445; 25%, 0.9460; 75%, 5.4733). Representative images are shown from n = 3 independent experiments. **b,** U2OS cells with different expression levels of Opto-G3BP1 were exposed to identical levels of blue light. Relative expression levels from top to bottom: 0.19, 0.32, 0.78 and 1 a.u. Representative images are shown from n = 3 independent experiments. **c,** U2OS cells stably expressing Opto-G3BP1 were pre-treated with cycloheximide (CHX) or ISRIB for 30 min and then exposed to 45 min of arsenite (NaAsO2) or 6 h of blue light stimulation, and immunostained with PABP antibody. **d,** Quantification of granule-positive cells from **c**. Data are shown as mean + s.e.m. from n = 3 independent experiments. **** *P* < 0.0001; ns, not significant by one-way ANOVA with Tukey post-test. **e,** Immunoblot showing phosphorylated eIF2α (p-eIF2α), eIF2α, and actin levels in cells treated with arsenite for 45 min, blue light for 6 h, or exposed to 42°C heat shock for 1 h. Scale bars, 10 μm in all micrographs.

Phase transitions are highly dependent on protein concentration, and we therefore hypothesized that the induction of OptoGranule assembly would be dependent on the local concentration of activated G3BP1, similar to the concentration-dependent formation of light-activated OptoDroplets^17^. To test this prediction, we controlled the local G3BP1 molecular concentration by modulating either the intensity of the activating blue light or the expression level of the Opto-G3BP1 construct. As predicted, we observed a strong positive correlation between blue light intensity and induction of OptoGranules (**Fig. 2a**) and, independently, a strong positive correlation between Opto-G3BP1 expression level and induction of OptoGranules (**Fig. 2b**). Thus, the OptoGranule system is highly tunable, a useful feature for a variety of studies.

We next examined the role of upstream events in OptoGranule formation and compared these to the cellular triggers associated with conventional stress granule assembly. Given that conventional stress granule formation is vitally linked to the disassembly of translating polysomes^16^, we tested whether polysome disassembly is required for OptoGranule formation. We determined that treatment with cycloheximide, which traps translating mRNAs within polysomes, blocked both the formation of arsenite-induced stress granules and the formation of light-induced OptoGranules (Fig. 2c,d), indicating that OptoGranule formation is dependent on polysome disassembly, further illustrating commonality with conventional stress granules. We next tested the role of eIF2α phosphorylation, which integrates stress granule formation downstream of a variety of stressors, such as arsenite and heat shock^16^. We used the small molecule ISRIB, which binds eIF2B and interrupts eIF2α-mediated translational control^18^. We found that formation of arsenite-induced stress granules was blocked by ISRIB, as previously documented^18^, whereas the formation of light-induced OptoGranules was unaffected by ISRIB treatment (Fig. 2c,d). Consistent with this finding, Western blotting also showed minimal phosphorylated eIF2α accompanying OptoGranule assembly (Fig. 2e). Thus, OptoGranule formation depends upon the recruitment of mRNPs from polysomes, but this assembly occurs downstream and independent of regulation by eIF2α.

We next examined the role of RNA in OptoGranule formation by using an RNA-binding assay to determine whether the Opto-G3BP1 fusion protein bound RNA similarly to G3BP1-GFP. Indeed, we found that Opto-G3BP1 bound bulk RNAs at levels comparable to G3BP1-GFP and this binding was unchanged by blue light activation (**Extended Data Fig. 3a**). Pursuing this result further, we modified the Opto-G3BP1 protein by deleting the RNA-binding domain of G3BP1 (Opto-G3BP1 ΔRBD) and tested the ability of this protein to form OptoGranules. As anticipated, Opto-G3BP1 ΔRBD did not form granules after activation with blue light, indicating that RNA binding is essential for the formation of OptoGranules (**Extended Data Fig. 3b,c**).

A subset of degenerative diseases, most notably ALS-FTD and inclusion body myopathy (IBM), are characterized by cytoplasmic inclusions composed of RNA-binding proteins and other constituents of RNA granules. A prominent feature of this end-stage cytoplasmic pathology is ubiquitinated and phosphorylated forms of TDP-43, although a host of other proteins co-localize with these pathological inclusions, including related RNA-binding proteins and ubiquitin-binding proteins such as p62/SQSTM1, UBQLN2, optineurin and VCP^19–23^. All of these proteins are components of stress granules and, moreover, mutations in many of these proteins are causative of ALS-FTD or IBM. Indeed, substantial evidence has accumulated implicating stress granules as the point of intersection for a wide array of insults that culminate in ALS-FTD or IBM, including mutations in RNA-binding proteins (e.g., hnRNPA1, TDP-43, TIA1, FUS), mutations in ubiquitin-binding proteins (VCP, UBQLN2, p62, OPTN), and pathological polydipeptides from repeat-expanded C9ORF72^1–13^.

Thus, it has been hypothesized that disturbances in the dynamics and material properties of membrane-less organelles such as stress granules may contribute to the initiation or progression of disease. A prediction of this hypothesis is that discrete disturbance in the dynamics and material properties of stress granules should be sufficient to cause cytotoxicity and recapitulate the pathognomonic features of these diseases. Until now, however, testing this hypothesis has been confounded by the requirement for exogenous stressors to induce chronic stress granules. To test this prediction in our system, we examined the consequences of chronic OptoGranule assembly. First, we examined the consequences of continuous blue light activation in cells expressing Opto-G3BP1 or Opto-Control. We found that chronic, persistent induction of OptoGranules resulted in progressive loss of cell viability accompanied by progressive loss of ATP levels (**Fig. 3a,b**). However, we noted that chronic exposure to blue light resulted in a moderate amount of cytotoxicity in cells expressing Opto-Control. Whereas cells expressing Opto-G3BP1 exhibited significantly greater loss of viability upon exposure to blue light than cells expressing Opto-Control, we sought to eliminate this background toxicity.

**Figure 3.**
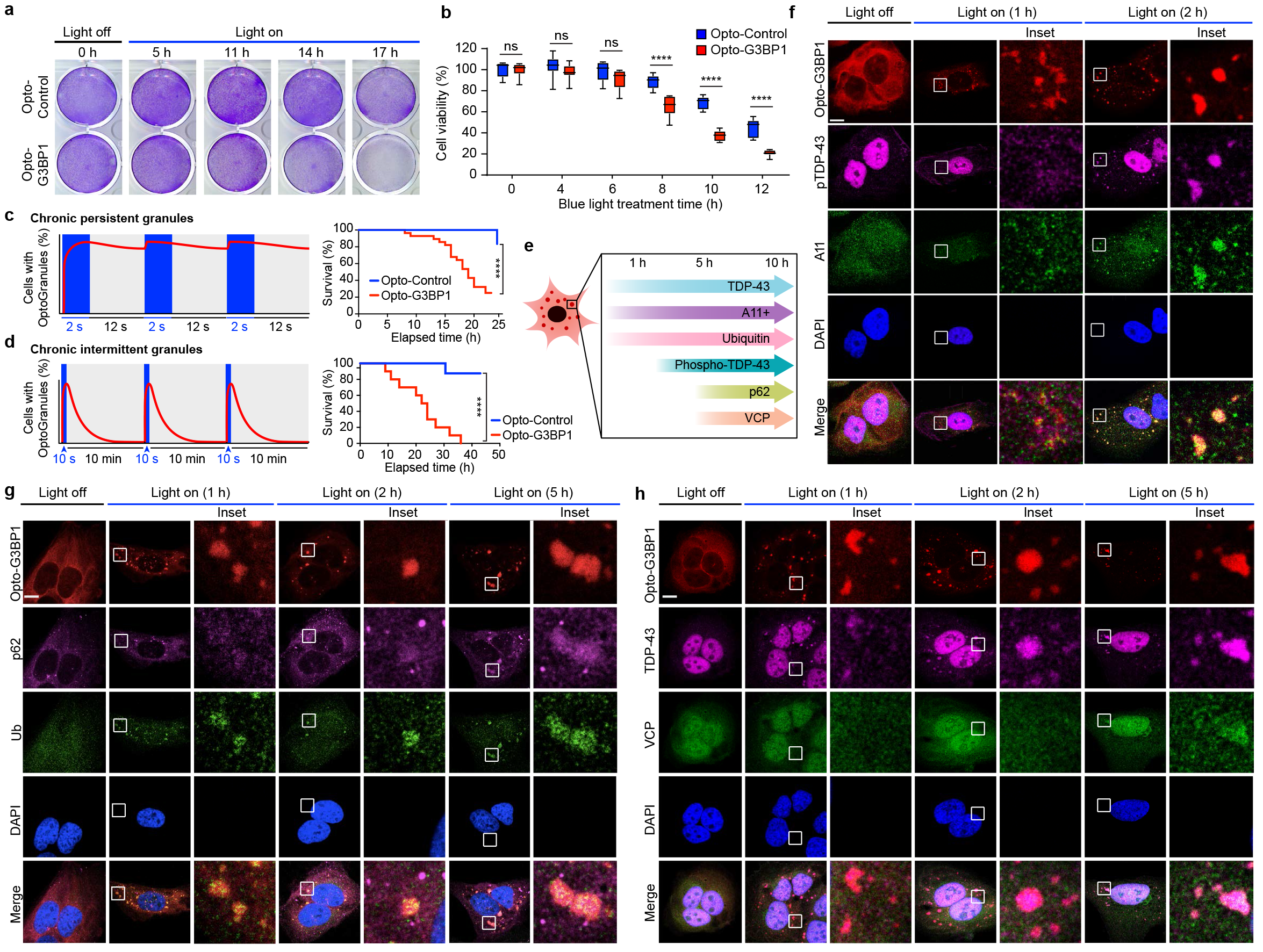
Persistent OptoGranules are cytotoxic and evolve to pathological inclusions. **a,b,** U2OS cells stably expressing Opto-Control or Opto-G3BP1 were stimulated with blue light and viability was assessed by crystal violet staining (**a**) or CellTiter-Glo 2.0 luminescence (**b**). Error bars represent s.d. from n = 9 biological replicates. \*\*\*\**P* < 0.0001.; ns, not significant by Student’s t test. **c,** U2OS cells stably expressing Opto-Control or Opto-G3BP1 were exposed to chronic persistent blue light stimulation with live cell imaging (left) and assessed for cell survival by counting living cells (right). n = 26 for Opto-Control and n = 28 for Opto-G3BP1. Data are shown from n = 3 independent experiments. \*\*\*\**P* < 0.0001 by log-rank (Mantel-Cox) test. **d,** U2OS cells stably expressing Opto-Control or Opto-G3BP1 were exposed to chronic intermittent blue light stimulation with live cell imaging (left) and assessed for cell survival by counting living cells (right). n = 7 for Opto-Control and n = 10 for Opto-G3BP1. \*\*\*\**P* < 0.0001 by log-rank (Mantel-Cox) test. **e**, Timeline of protein accumulation in OptoGranules. **f-h,** U2OS cells stably expressing Opto-G3BP1 were stimulated with blue light for indicated times and co-immunostained with p-TDP-43 and A11 antibodies (**f**), p62 and ubiquitin antibodies (**g**), or VCP and TDP-43 antibodies (**h**). Images in **f-h** are representative of n = 3 independent experiments. Scale bars, 10 μm in all micrographs.

Thus, we used live, confocal based imaging to monitor cell viability in real time during laser-induced OptoGranule induction at 445 nm. We first used a paradigm consisting of 2-second blue light pulses alternating with 12 seconds of rest, which drove robust OptoGranule assembly but left insufficient time for granules to disassemble prior to the next light pulse, resulting in persistent OptoGranule assembly (**Fig. 3c**). Interestingly, persistent OptoGranule assembly under these conditions (2 sec on, 12 sec off) resulted in significant loss of viability in cells expressing Opto-G3BP1 and greatly reduced toxicity in cells expressing Opto-Control (**Fig. 3c**). Pursuing this further, we established a paradigm that even further minimized blue light exposure, using a 10 seconds blue light pulse, followed by 10 minutes of rest, which was sufficient to initiate robust assembly of OptoGranules that were able to fully disassemble prior to the next light pulse (**Fig. 3d**). This paradigm of chronic, intermittent OptoGranule assembly, which may more closely reflect physiological, chronic, intermittent stress, eliminated background toxicity due to blue light exposure and revealed significant toxicity in cells expressing Opto-G3BP1 compared to cells expressing Opto-Control (**Fig. 3d**). Thus, we conclude that chronic persistent or chronic intermittent stress granule assembly is intrinsically cytotoxic, independent of exogenous stressors.

Disease pathology in tissue from patients with ALS and FTD is marked by deposits of ubiquitin, ubiquitin-binding proteins, and TDP-43 that is cleaved and abnormally phosphorylated at Ser409/410^24^. Newly formed OptoGranules were easily distinguished from the pathology present in late-stage ALS and FTD. Although OptoGranules were initially immunopositive for TDP-43 (as are conventional arsenite-induced stress granules), they were immunonegative for phospho-TDP-43, ubiquitin and ubiquitin-binding proteins. OptoGranules were, however, immunopositive for staining by anti-A11, a conformation-specific antibody that recognizes amyloid oligomer, a feature also shared by conventional stress granules (**Fig. 3e,f** and **Extended Data Fig. 4a**). The presence of A11 immunopositivity in newly formed stress granules suggests that non-pathological amyloid oligomers are present in the mRNPs recruited to these structures, perhaps arising from the prion-like low complexity domains of RNA-binding proteins coating these mRNPs. While these are presumably physiological amyloids, it is conceivable that their close packing in the condensed liquid state of persistent stress granules risks seeding the assembly of pathological amyloids, particularly for proteins like TDP-43 that can adopt highly stable structures.

Remarkably, the characteristics of OptoGranules changed during chronic assembly. Specifically, we observed that after approximately two hours of OptoGranule assembly, there was a significant increase in immunopositivity using two distinct anti-ubiquitin antibodies and two distinct anti-phospho-TDP-43 antibodies (**Fig. 3f, Extended Data Fig. 4b,c**). The anti-phospho-TDP-43 antibodies specifically recognize phosphorylation of TDP-43 at residues Ser409/410, a pathological signature specific to a spectrum of sporadic and familial forms of TDP-43 proteinopathies, including ALS-FTD^24^. Moreover, after approximately five hours of chronic OptoGranule assembly, we observed a significant increase in immunopositivity using antibodies to the ubiquitin-binding proteins p62/SQSTM1 and VCP, illustrating further evolution of these structures (**Fig. 3g,h and Extended Data Fig. 4d**). Thus, not only does chronic OptoGranule assembly cause a loss of cell viability, but cell death is preceded by the evolution of OptoGranules into cytoplasmic inclusions that recapitulate features that are pathognomonic for ALS-FTD.

We next examined the disease relevance of these findings in a neuronal context by generating human induced pluripotent stem cell (iPSC)-derived neurons. In response to arsenite or heat shock stresses, these iPSC-derived neurons assembled conventional stress granules that were positive for TIA1 and TDP-43, indicating that they are suitable for examining the consequences of chronic stress granules (**Extended Data Fig. 5a**). Next, we introduced Opto-G3BP1 into differentiated neurons (mRuby-tagged Opto-G3BP1) (**Fig. 4a**) or by integration into iPSCs prior to differentiation (Doxycycline inducible mCherry-tagged Opto-G3BP1) (**Extended Data Fig. 5c**). In mRuby-Opto-G3BP1-expressing neurons, blue light activation induced the assembly of OptoGranules indistinguishable from those observed in U2OS cells (**Fig. 4b**, **Extended Data Fig. 5b and Supplementary Videos 4, 5**). Chronic induction of OptoGranules following transient introduction of mRuby-Opto-G3BP1 resulted in progressive loss of neuronal viability (**Fig. 4c**) and the formation of neuronal cytoplasmic inclusions that were immunopositive for TDP-43 and A11 (**Fig. 4d-f**) with time-dependent immunopositivity for phosphorylated TDP-43, ubiquitin and p62/SQSTM1 (**Fig. 4g,h**). Lastly, we examined neurons derived from iPSCs stably expressing inducible Opto-G3BP1 (**Extended Data Fig 5c**). During differentiation into neurons, Opto-G3BP1 expression was also induced by adding doxycycline and remained diffuse until activation with blue light, whereupon these neurons assembled dynamic OptoGranules that evolved into neuronal cytoplasmic inclusions with chronic stimulation (**Extended Data Fig. 5d-h**). Thus, chronic OptoGranule induction recapitulates the evolution of ALS-FTD pathology and neurotoxicity in human iPSC-derived neurons.

**Figure 4.**
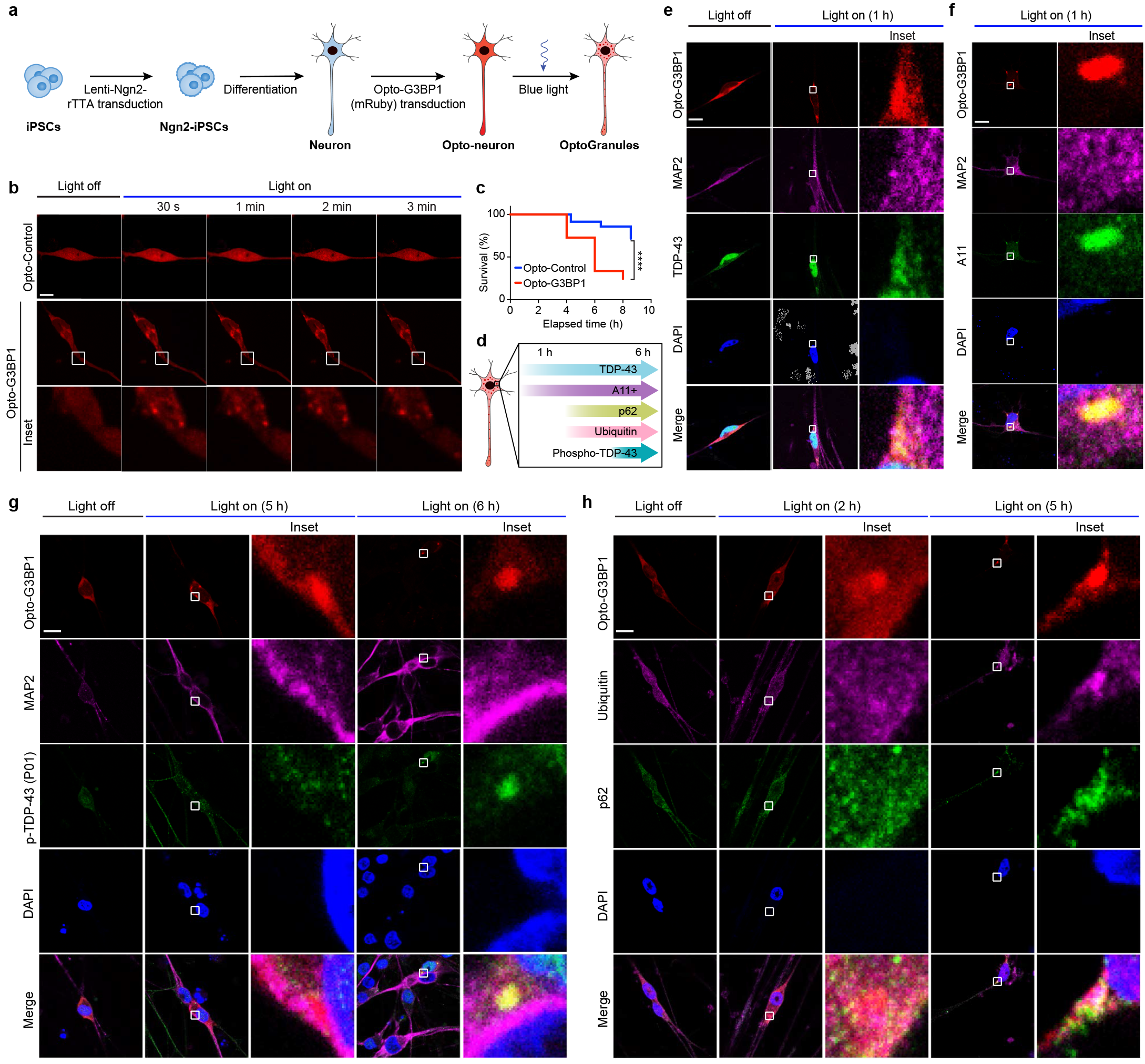
Persistent OptoGranules are cytotoxic and evolve to pathological inclusions in human iPS cell-derived neurons. **a,** Schematic of Opto-neuron generation. **b,** iPS cell-derived neurons expressing Opto-Control (mRuby) or Opto-G3BP1 (mRuby) were exposed to identical blue light stimulation. Representative images are shown from n = 3 independent experiments. **c,** iPS cell-derived neurons expressing Opto-Control or Opto-G3BP1 were exposed to chronic persistent stimulation and survival was assessed by counting living cells. n = 35 cells for Opto-Control and n = 34 cells for Opto-G3BP1. Data are representative of n = 3 independent experiments. \*\*\*\**P* < 0.0001 by log-rank (Mantel-Cox) test. **d,** Timeline of pathological protein accumulation in OptoGranules in iPS cell-derived neurons. **e-h,** iPS cell-derived neurons expressing Opto-G3BP1 were stimulated with blue light for indicated times and co-immunostained with MAP2 and TDP-43 antibodies (**e**), MAP2 and A11 antibodies (**f**), MAP2 and p-TDP-43 (P01) antibodies (**g**), or p62 and ubiquitin antibodies (**h**). Images in **e-h** are representative of n = 3 independent experiments. Scale bars, 10 μm in all micrographs.

Among the many membrane-less organelles that arise through phase transitions, stress granules have drawn the most attention from the ALS-FTD field because of their cytoplasmic location, which matches the location of pathological deposits in ALS-FTD, and the many disease-associated proteins that are components of stress granules. However, we must emphatically note that phase separation-mediated assembly, dynamics, and material properties of stress granules must be viewed within the context of a larger cellular network of membrane-less organelles, which include a wide variety of nuclear and cytoplasmic RNA granules. Indeed, membrane-less organelles are now recognized as functionally relevant biomolecular condensates that underlie different segregated biochemistries within a single cell^13^. Furthermore, their material properties (e.g., assembly/disassembly rates, mobility, viscosity) likely influence these functions; indeed, the data presented here supports the burgeoning hypothesis that ALS-FTD arises from disturbances in the dynamics and material properties of membrane-less organelles, with devastating consequences over time.

Extending this hypothesis, we speculate that disease may reflect simultaneous pathological disturbance of multiple membrane-less organelles that arises by derangement of a network of multiple, independent phases. These interconnections likely reflect communication across different types of membrane-less compartments based on rapid, dynamic exchange of macromolecules (e.g., RNA and RNA-binding proteins) and small molecules that act as vehicles to communicate material states throughout the network. An example of this is seen in the recent report that perturbation of one phase separated compartment (stress granules) alters the properties and function of a distinct phase separated structure (the nuclear pore)^25^. With such a system-wide regulation, primary disturbances in the material properties of one node of the network (e.g., stress granules) may lead to secondary disturbances that are propagated throughout the entire network of membrane-less organelles.

## Methods

### Cell culture and transfection

U2OS cells were purchased from ATCC (HTB-96). U2OS cells were cultured in Dulbecco’s modified Eagle’s medium (HyClone) supplemented with 10% fetal bovine serum (Hyclone SH30071.03 and SH30396.03), 1X GlutaMAX (Thermo Fisher Scientific 35050061), 50 U/ml penicillin, and 50 μg/ml streptomycin (Gibco 15140-122), and maintained at 37°C in a humidified incubator with 5% CO_2_. FuGENE 6 (Promega E2691) was used for transient transfections per the manufacturer’s instructions. *G3BP* KO cells have been previously described^1^. U2OS cells stably expressing G3BP1-GFP have been previously described^2^. Cells were regularly checked for mycoplasma by DAPI staining.

### Plasmids

DNA fragments encoding human G3BP1 and TIA1 were PCR-amplified from G3BP1 (DNASU HsCD00042033) and pEGFP-TIA1^3^, respectively. FUS and TDP-43 were PCR-amplified from cDNA. The pCRY2PHR-mCherry backbone was PCR-amplified from pCRY2PHR-mCherryN1 (Opto-Control; Addgene 26866). DNA fragments encoding G3BP1, TIA1, TDP-43, and FUS were inserted into pCRY2PHR-mCherryN1 backbone using NEBuilder HiFi DNA Assembly Master Mix kit (NEB E2621). To create Opto-G3BP2, DNA fragments encoding G3BP2 were amplified from cDNA and inserted into pCMV-CRY2-mCherry at XhoI and BamHI using NEBuilder HiFi DNA Assembly Master Mix. Mammalian codon-optimized pCRY2PHR-mCherry was PCR-amplified from pCMV-CRY2-mCherry (Addgene 58368). mRuby3 was PCR-amplified from pKanCMV-mClover3-mRuby3 (Addgene 74252). Opto-G3BP1 (mRuby) was assembled from codon-optimized pCRY2PHR-mCherry, mRuby3, and G3BP1 DNA using NEBuilder HiFi DNA Assembly Master Mix kit. Opto-G3BP1 (mRuby) lentiviral plasmids were constructed by inserting PCR-amplified CMV-promoted CRY2-mRuby-G3BP1 (dNTF2) into PspXI and EcoRI linearized cloning backbone phND2-N174 (Addgene 31822) using NEBuilder HiFi DNA Assembly Master Mix kit. Dox-Opto-G3BP1 (mCherry) lentiviral plasmids were constructed by inserting PCR-amplified Opto-Control and Opto-G3BP1 into EcoRI-digested cloning backbone pTight-hND2-N106 (Addgene 31875) using NEBuilder HiFi DNA Assembly Master Mix kit. Truncations were introduced using Q5 site-directed mutagenesis (NEB E0054). G3BP1-GFP constructs have been previously described^4^. All constructs were confirmed by sequencing.

### Drugs and heat shock treatments

ISRIB (200 nM; Sigma SML0843) and cycloheximide (100 μg/ml; Sigma C4859) treatment was performed for 30 min before adding sodium arsenite (0.5 mM; Sigma 35000-1L-R) or blue light. For sodium arsenite treatment, medium was changed to medium containing 0.5 mM sodium arsenite for 45 min. For heat shock treatment, cells were transferred to a 42°C humidified incubator with 5% CO_2_ for 1 h.

### Lentivirus production

Lenti-X 293T cells (293LE; Clontech 632180) were transfected at 80–90% confluency with viral vectors containing genes of interest and viral packaging plasmids psPAX2 (Addgene 12260) and pMD2.G (Addgene 12259) using polyethylenimine (Polysciences 24765-2). The medium was changed 24 h after transfection. Viral supernatants were harvested at 48 h after transfection, filtered with 0.45 μM filters, and centrifuged at 100,000 × g at 4°C for 1.5 h. Ultracentrifugation was carried out through a 20% (w/v in PBS) sucrose cushion at 100,000 × g at 4°C for 1.5 h. Pellets were resuspended in 100 μl DMEM + 10% FBS and stored at −80°C.

### Stable cell lines

Opto-Control (mCherry) or Opto-G3BP1 (mCherry) constructs were cotransfected with linear hygromycin marker (Clontech 631625) into U2OS cells using FuGENE 6 (Promega). 48 h after transfection, 200 μg/ml hygromycin B (Thermo Fisher Scientific 10687010) was added to culture media for selection. mCherry-positive cells were selected using cell sorting to produce Opto-Control (mCherry) or Opto-G3BP1 (mCherry) stable cell lines. Filtered Opto-Control (mRuby) or Opto-G3BP1 (mRuby) viral supernatants and 8 μg/ml polybrene (Sigma H9268) were added to U2OS cells at ~50% confluency in 10-cm plates. mRuby-positive cells were selected using cell sorting to produce Opto-Control (mRuby) or Opto-G3BP1 (mRuby) stable cell lines.

### iPSC neuron differentiation

iPSC neurons were generated as described previously^5^ with modifications. iPSCs ((Re)Building a Kidney Q-2D6W) were dissociated with Gentle Cell Dissociation Reagent (Stemcell Technologies 07174) and 300,000 iPSCs were seeded into one Matrigel (Corning 354277)-coated well of a six-well plate in mTeSR medium (Stemcell Technologies 85850) containing 10 μM ROCK inhibitor (Stemcell Technologies 72302). The next day, the medium was changed to mTeSR medium.

To generate Opto-neurons (mRuby), lentiviruses encoding Ngn2 and rTTA were added to the medium at MOI = 4, respectively, in the presence of hexadimethrine bromide (4 μg/ml; Sigma-Aldrich H9268) and the medium was changed 24 h after transduction. When transduced iPSCs reached 75% confluency, 1 μg/ml of doxycycline hyclate (Sigma-Aldrich D9891) was added to mTeSR medium to induce Ngn2 expression. At day 2 of induction, iPSCs were dissociated with Gentle Cell Dissociation Reagent and 150,000 cells were seeded onto coverslips in one well of a 24-well plate or 4-well Nunc Lab-Tek chambered coverglass (Thermo Fisher Scientific 155382) coated with Poly-L-ornithine/laminin/fibronectin (Sigma-Aldrich P4957; Sigma-Aldrich L2020; Sigma-Aldrich F4759^6^), and cultured in BrainPhys neuronal medium (Stemcell Technologies 05790) containing 1 × N2 (Thermo Fisher Scientific 17502048), 1 × B27 (Thermo Fisher Scientific 12587010), 20 ng/ml BDNF (Peprotech 450-02), 20 ng/ml GDNF (Peptrotech 450-10), 500 μg/ml Dibutyryl cyclic-AMP (Sigma-Aldrich D0627), 200 nM L-ascorbic acid (Sigma-Aldrich A0278), 1 μg/ml natural mouse laminin (Thermo Fisher Scientific 23017-015), 1 μg/ml DOX, and 1 μg/ml puromycin (Thermo Fisher Scientific A1113803). Opto-Control (mRuby) or Opto-G3BP1 (mRuby) lentiviruses and 4 μg/ml hexadimethrine bromide (Sigma H9268) were added to iPSC neurons at 3-5 DIV. Media was changed approximately 12 h after transduction and then half-changed every other day until the assay was performed.

To generate doxycycline inducible Opto-neurons (mCherry), lentiviruses encoding Ngn2, rTTA, and Dox-Opto-Control or Dox-Opto-G3BP1 were added to the medium at ~MOI = 4, respectively, in the presence of hexadimethrine bromide (4 μg/ml), and the medium was changed 24 h after transduction. iPSCs were dissociated with Gentle Cell Dissociation Reagent and 150,000 cells were seeded into coverslips in one well of a 24-well plate or 4-well Nunc Lab-Tek chambered coverglass coated with Matrigel/Poly-L-ornithine/laminin/fibronectin and cultured in BrainPhys neuronal medium for 7 days. iPSC neuron cultures were maintained in BrainPhys neuronal medium and half-changed every other day until the assay was performed.

### Blue light LED treatment

Cells (30-60% confluency) were transferred into blue light illumination at ~2 mW/cm^2^ using custom-made LED arrays in a humidified incubator with 5% CO_2_ with blue light LED array. Custom-made LED arrays were arranged with a flexible LED strip light (Ustellar). The light intensity of LED arrays was measured by a power meter (ThorLabs S170C).

### Live cell imaging

All live-cell imaging experiments were performed using a Marianas 2 spinning disk confocal imaging system except overnight images of cell viability assays (described below). Images were acquired using a 63×/1.4 Plan Apochromat objective. Cells were plated in 4-well Nunc Lab-Tek chambered coverglass (Thermo Fisher Scientific 155382). Before imaging, the medium was changed to FluoroBrite DMEM medium (Thermo Fisher Scientific A1896701) with 10% fetal bovine serum and 1X GlutaMAX. During imaging, cells were maintained at 37°C with an environmental control chamber. Definite focus was used during the live-cell imaging. For one-time photoactivation, indicated cells were initially photoactivated by a 5-ms pulse of 488-nm laser illumination at 55% of maximum laser power, then imaged every 1 s thereafter with a 561-nm laser. For repeated activation, images were taken with 488-nm and 561-nm channels with 100 ms exposure time. Images were analyzed with SlideBook 6 software.

### Western blotting

Cells were collected using PBS and lysed for 10 min on ice using RIPA buffer (25 mM Tris-HCl (pH 7.6), 150 mM NaCl, 1% NP-40, 1% sodium deoxycholate, 0.1% SDS; Pierce, 89901) supplemented with proteinase inhibitor cocktail (Roche 1186153001) and PhosSTOP (Roche 04906845001). Samples were centrifuged for 20 min at 4°C at 14,000 rpm. 4X NuPAGE LDS sample buffer (Thermo Fisher Scientific NP0008) was added to the supernatant and samples were boiled for 5 min. Samples were run in 4-12% NuPAGE Bis-Tris gels (Invitrogen) and transferred to nitrocellulose membranes using an iBlot 2 transfer device (Thermo Fisher Scientific). Membranes were blocked with Odyssey blocking buffer (LI-COR) and then incubated with primary antibodies. Following incubation with dye-labeled secondary antibodies, signals were visualized using an Odyssey Fc imaging system (LI-COR). Primary western blot antibodies were anti-β-actin (Santa Cruz Biotechnology sc-1616), anti-eIF2α (Santa Cruz Biotechnology sc-133132), anti-phospho-eIF2α (Cell Signaling 3597S), anti-mCherry (Abcam 167453), and anti-G3BP1 (BD biosciences 6111126). Secondary western blot antibodies were IRDye 800CW donkey anti-mouse, IRDye 800CW donkey anti-goat, and IRDye 680RD donkey anti-rabbit (LI-COR 926-32212, 926-32214, and 926-68073 respectively) and used at a dilution of 1:15,000.

### Immunofluorescence

Cells were grown in 8-well chamber slides (Millipore). Following the indicated stimulation, cells were fixed with 4% paraformaldehyde (Electron Microscopy Science) in PBS for 10 min at room temperature, permeabilized with 0.2% Triton X-100 in PBS for 10 min at room temperature, and then blocked with 10% normal goat serum (Life Technologies 50062) or 5% BSA for 1 h at room temperature. Samples were incubated with primary antibodies in blocking buffer overnight at 4°C. Samples were then washed three times with PBS and incubated with secondary antibody for 1 h at room temperature. Primary antibodies were anti-PABP (Abcam ab21060), anti-G3BP1 (BD Biosciences 6111126), anti-eIF4G (Santa Cruz Biotechnology sc-11373), anti-TDP-43 (Proteintech 12892-1-AP), anti-phospho-TDP-43 (M01) (Cosmo Bio CO TIP-PTD-MO1), anti-phospho-TDP-43 (P01) (Cosmo Bio CO TIP-PTD-PO1), anti-VCP (BD Biosciences 612183), anti-amyloid-oligomer A11 (Thermo Fisher Scientific AHB0052), anti-Ubiquitin (Dako, Z0458), anti-Ubiquitin (Santa Cruz Biotechnology sc-8017), anti-p62 (Abcam 56416), anti-MAP2 (Sigma M9942), anti-TIA1 (Santa Cruz Biotechnology sc-1751), anti-TIAR (BD Biosciences 610352), anti-eIF3η (Santa Cruz Biotechnology sc-16377), anti-ataxin 2 (Proteintech 21776-1-AP), and anti-GLE1 (Abcam 96007). Secondary antibodies were Alexa Fluor 488/555/647 (Life Technologies). For microscopic imaging, slides were mounted with ProLong Gold Antifade Mountant with DAPI (Invitrogen). Images were captured using a Leica TCS SP8 STED 3X confocal microscope with a 63x oil objective.

### Fluorescence recovery after photobleaching

Cells were first activated by dual imaging every 1 s for 5 min with 55% 488-nm laser power and 100-ms exposure time to initiate granule formation. Opto-G3BP1 or G3BP1-GFP-positive stress granules were then photobleached and mCherry or GFP signal intensity was measured before and after photobleaching.

### Fluorescent *in situ* hybridization

Cells were fixed with 4% paraformaldehyde at room temperature for 10 min and then washed twice with PBS. 70% (v/v) EtOH was then added and cells were stored at 4°C overnight. Cells were then washed twice with wash buffer (2X SSC with 10% formamide in RNase-free water). Following aspiration of the wash buffer, cells were incubated with hybridization buffer (2x SSC, 10% v/v deionized formamide, 10% (w/v) dextran sulfate, 2 mM vanadyl ribonucleoside complex, 1 mg/ml yeast tRNA (Ambion AM7119), 0.005% BSA (Ambion AM2616) with 1 ng/μl 5′ labeled FAM-oligo(dT20) probes (Genelink 26-4620-02) at 37°C overnight. Cells were then washed 3 times with pre-warmed wash buffer at 37°C.

### RNA binding assay

The RNA binding assay was adapted from a previous report^7^. Cells were rinsed with PBS and UV cross-linked on ice using a Stratagene UV cross-linker at 600 mJ/cm^2^. Cells were lysed in 0.5 ml/dish lysis buffer (25 mM Tris-HCl pH 7.5, 137 mM NaCl, 1% Triton X-100, 2 mM EDTA, 1x protease inhibitor (Roche Diagnostics 11836145001)) for 10 min on ice. One unit of RQ1 DNase (Promega M6101) was added to the cell lysates and incubated at 37°C for 5 min at 1000 rpm in a Thermomixer (Eppendorf). Two units of RNase I (Life Technologies AM2294) was added and lysates were further incubated for 3 min at 37°C under 1000 rpm agitation in the Thermomixer. Supernatants were collected after 21,000 × g centrifugation at 4°C for 10 min. 10 μl RFP-Trap (Chromotek rtma-20) or GFP-Trap beads (Chromotek gtma-20) were incubated with the supernatant fraction at 4°C overnight. The beads were washed twice with lysis buffer, twice with 1M NaCl, and twice with lysis buffer again. Beads were further suspended in 100 μl 10 mM Tris-HCl pH 7.5 and treated with 2 units of calf intestinal alkaline phosphatase (Promega M182A) at 37°C for 15 min at 1000 rpm in the Thermomixer. Beads were washed with lysis buffer twice and RNA labeling was performed with RNA 3′ End Biotinylation Kit (Pierce 20160). After washing with lysis buffer twice, beads were boiled in 1x LDS sample buffer (Life Technologies NP007) with 100 mM DTT. Protein-RNA complexes were separated by 4-12% NuPAGE Bis-Tris gels, transferred to nitrocellulose membranes, and blotted with IRDye 680LT Streptavidin (LI-COR 926-68031).

### Crystal violet assay

Cells were seeded at ~20% confluency in 6-well plates and grown for 24 h before exposure to LED blue light. At the indicated treatment time, media was aspirated and replaced with staining solution (0.05% (w/v) crystal violet, 1% formaldehyde, 1% methanol in 1X PBS) for 20 min at room temperature followed by 3 washes with water.

### CellTiter-Glo 2.0 cell viability assay

This assay determines the number of viable cells by measuring ATP, which indicates the presence of metabolically active cells. Cells were seeded at 4-5 × 10^3^ cells/well in 96-well plates one day before exposure to blue light LED. Following blue light exposure, cells were measured using the CellTiter-Glo 2.0 assay kit (Promega G9242) per the manufacturer’s instructions.

### Neuron viability imaging

Opto-Control and Opto-G3BP1 mRuby neurons were imaged on a DMI8 Widefield Microscope (Leica) with a 20x Plan Apo 0.80NA air objective using LAS X 3.4.2.18368 software (Leica). By saving the stage positions, a tilescan capture was taken in the same location every 2 h using the 561-nm filter at 600-ms exposure. Between imaging, neurons were placed in the blue light LED incubator until the next time point. Stitching was performed in LAS X, with each merged image totaling an area ~37.82 mm^2^.

### Overnight live cell imaging

Overnight live cell imaging experiments were performed with an Opterra II Swept Field confocal microscope (Bruker) using Prairie View 5.4 Software. Opto-Control and Opto-G3BP1 cells were plated in the middle two wells of a 4-well Lab-Tek chambered coverglass (Nunc) at ~20% confluency the day prior to imaging. Immediately before imaging, the medium was changed to FluoroBrite DMEM medium supplemented with 10% fetal bovine serum and 1X GlutaMAX. During imaging, cells were maintained at 37°C and supplied with 5% CO_2_ using a Bold Line Cage Incubator (Okolabs) and an objective heater (Bioptechs). Imaging was performed using a 60x Plan Apo 1.40NA oil objective and Perfect Focus (Nikon) was engaged for the duration of the capture. Continuous activation data was acquired with a script made in Prairie View. The script was set to image the 561-nm channel with 100-ms exposure at 80 power in a multipoint capture once, followed by imaging the 445-nm channel with 2000-ms exposure at 200 power in a multipoint capture five times. This script was repeated continually for the duration of the experiment. Three fields of Opto-Control and Opto-G3BP1 cells each with similar expression levels were chosen per experiment. Analysis was performed using ImageJ.

### Statistical analysis

*P* > 0.05 was considered not significant. \**P* ≤ 0.05, \*\**P* < 0.01, and \*\*\**P* < 0.001 by one-tailed Student’s t test, one-way ANOVA or Log-rank (Mantel-Cox) test. Statistical analysis was performed in GraphPad Prism or Excel.

## Data availability

The data that support the findings of this study are available from the corresponding author on reasonable request.

## Author Contributions

J.P.T. conceived and supervised the project. P.Z., B.F., P.Y., J.T. and J.M. designed and performed the experiments and data analyses. P.Z, H.J.K. and J.P.T. wrote the manuscript.

## Acknowledgements

We thank Natalia Nedelsky for editorial assistance. We thank Anderson Kanagaraj for assistance with DNA construct preparation. We also thank Dr. Gitler for kindly providing with phospho-TDP-43 antibodies. This work was supported by grants HHMI, NIH grant R35NS097974, and St. Jude Research Collaborative Membrane-less Organelles to J.P.T. J.P.T. is a consultant for Third Rock Ventures.

---

**Extended Data Figure 1.**
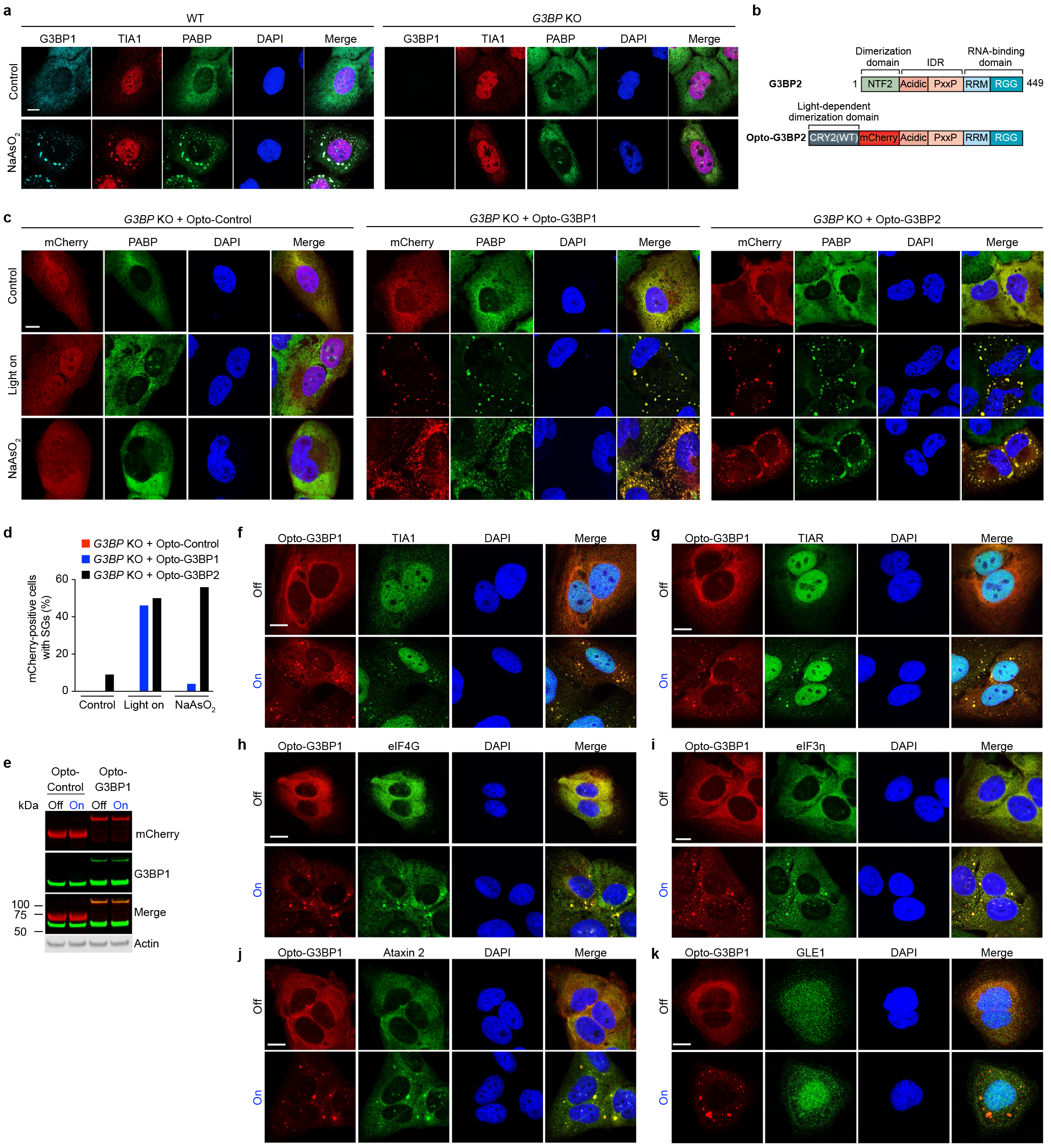
OptoGranules are light-inducible stress granules. **a,** WT U2OS cells and *G3BP1/2* double KO cells with or without sodium arsenite (NaAsO_2_) treatment were immunostained for G3BP1 as well as stress granule markers TIA1 and PABP. Representative images are shown from n = 3 independent experiments. **b,** Schematic diagram of Opto-G3BP2 construct. **c,** *G3BP1/2* double KO cells transiently transfected with Opto-Control, Opto-G3BP1, or Opto-G3BP2 constructs were treated with sodium arsenite (NaAsO_2_) for 45 min or exposed to 6 h of blue light and immunostained for the stress granule marker PABP. Representative images are shown from n = 3 independent experiments. **d,** Quantification of PABP-positive stress granules in **c**. **e,** Immunoblot showing expression level of mCherry (red) and G3BP1 (green) in U2OS cells stably expressing Opto-Control or Opto-G3BP1 with or without blue light stimulation. Representative blot is shown from n = 3 independent experiments. **f-k,** U2OS cells stably expressing Opto-G3BP1 with or without blue light stimulation were immunostained with antibodies against TIA1 (**f**), TIAR (**g**), eIF4G (**h**), eIF3n (**i**), ataxin 2 (**j**), or GLE1 (**k**). Representative images are shown from n = 3 independent experiments. Scale bars, 10μm in all micrographs.

**Extended Data Figure 2.**
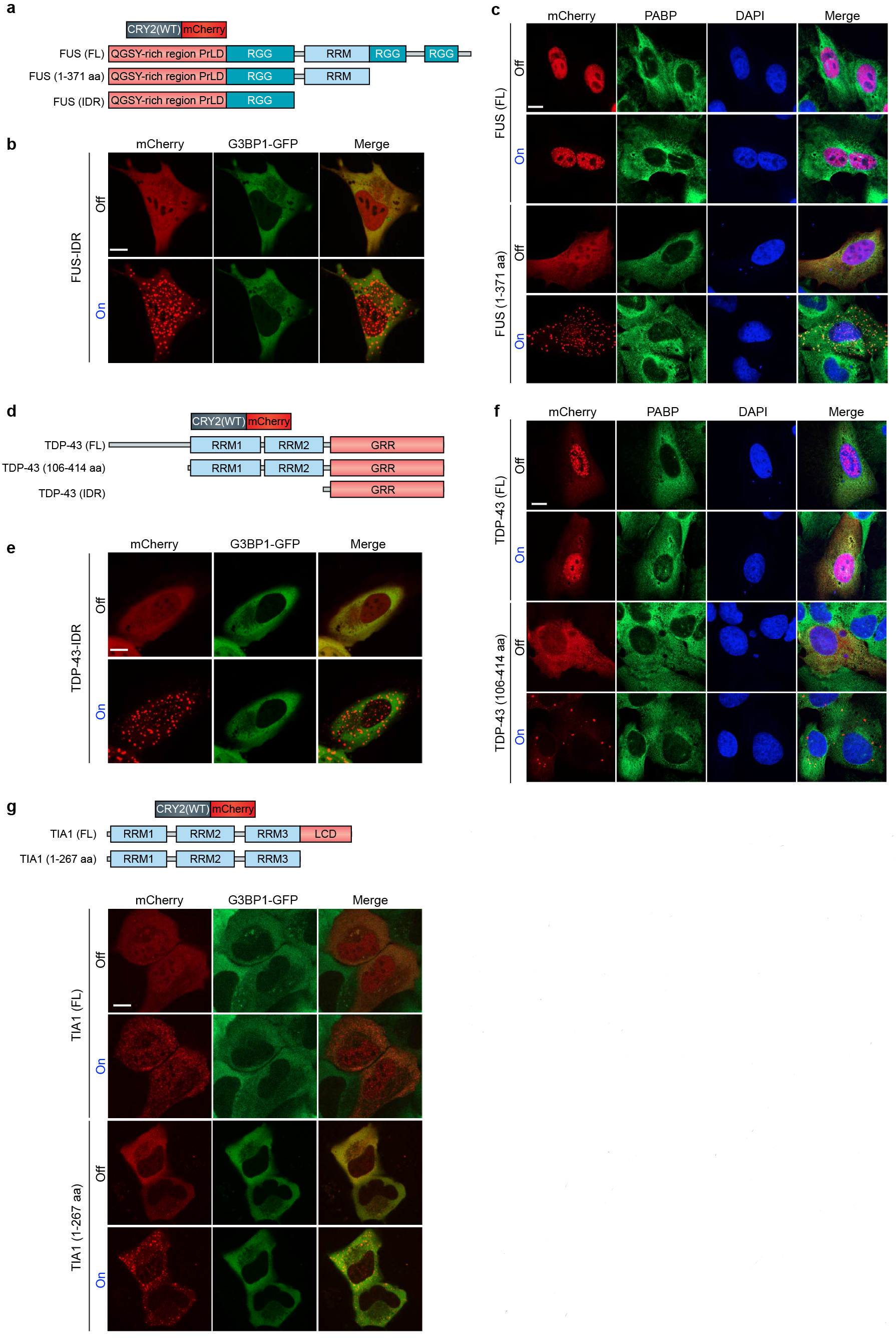
OptoDroplets are not stress granules. **a,** Schematic diagram of Opto-FUS constructs. **b,** U2OS cells were transiently co-transfected with Opto-FUS-IDR and stress granule marker G3BP1-GFP, and imaged before and after blue light stimulation. Representative images are shown from n = 3 independent experiments. **c,** U2OS cells were transiently transfected with Opto-FUS (full-length, FL) or Opto-FUS (1-371 aa) and imaged before and after blue light stimulation by immunostaining with stress granule marker PABP. Representative images are shown from n = 3 independent experiments. **d,** Schematic diagram of Opto-TDP-43 constructs. **e,** U2OS cells were transiently co-transfected with Opto-TDP-43-IDR and stress granule marker G3BP1-GFP, and imaged before and after blue light stimulation. Representative images are shown from n = 3 independent experiments. **f,** U2OS cells were transiently transfected with Opto-TDP-43 (FL) or Opto-TDP-43 (106-414 aa) and imaged before and after blue light stimulation by immunostaining with stress granule marker PABP. Representative images are shown from n = 3 independent experiments. **g,** Top: Schematic diagram of Opto-TIA1constructs. Bottom: U2OS cells were transiently co-transfected with Opto-TIA1 (full-length) or Opto-TIA1 (1-267 aa) and stress granule marker G3BP1-GFP, and imaged before and after blue light stimulation. Representative images are shown from n = 3 independent experiments. Scale bars, 10 μm in all micrographs.

**Extended Data Figure 3.**
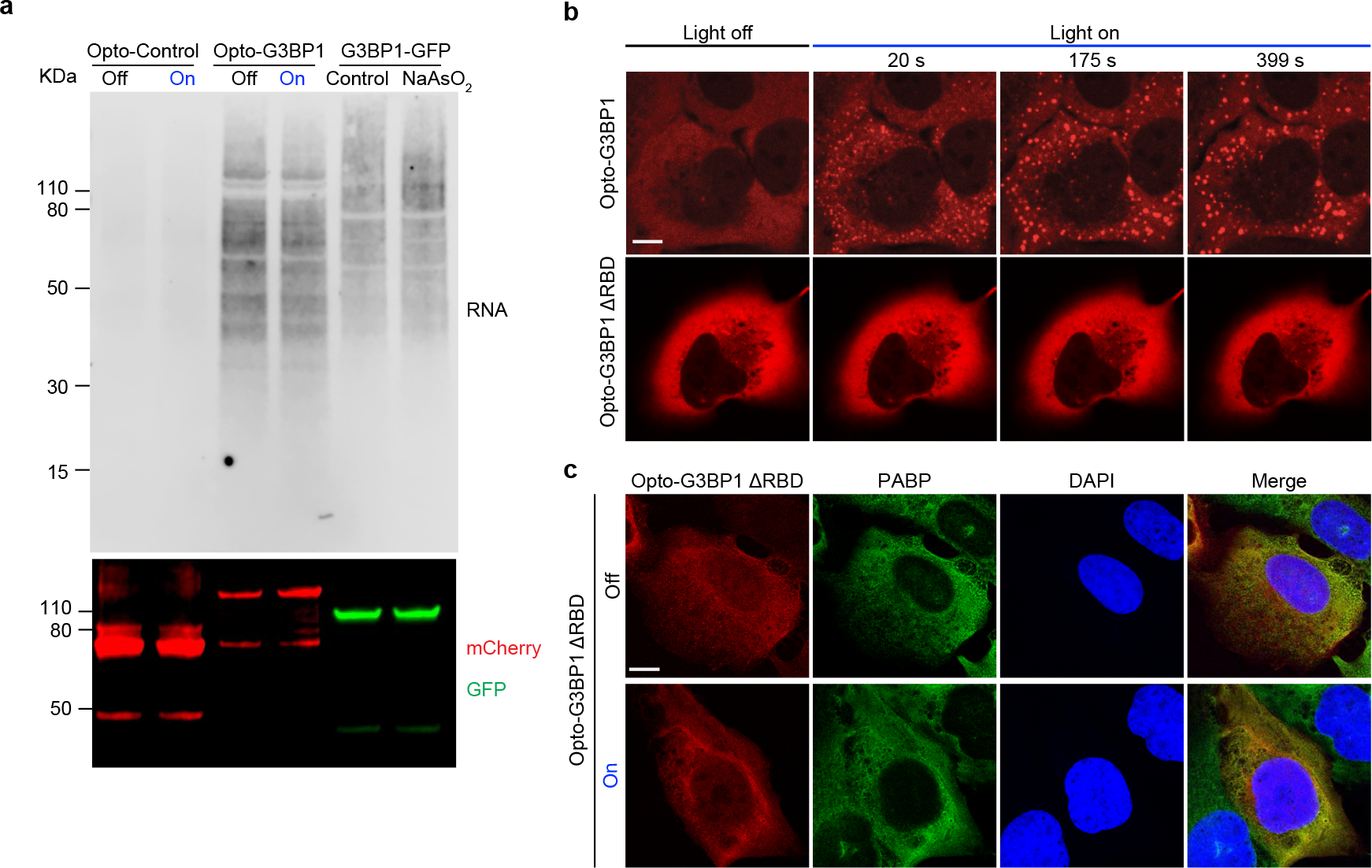
OptoGranule formation is dependent on RNA binding. **a,** U2OS cells stably expressing Opto-Control, Opto-G3BP1, or G3BP1-GFP were treated with blue light for 6 h or sodium arsenite for 30 min and then blotted with IRDye 680LT Streptavidin or mCherry after RNA-binding assay. **b,** U2OS cells transiently transfected with Opto-G3BP1 or Opto-G3BP1-ΔRBD were stimulated with blue light for indicated times. Representative images are shown from n = 3 independent experiments. **c,** U2OS cells transiently transfected with Opto-G3BP1-ΔRBD were exposed to blue light stimulation for 6 h and immunostained for the stress granule marker PABP. Representative images are shown from n = 3 independent experiments. Scale bars, 10 μm in all micrographs.

**Extended Data Figure 4.**
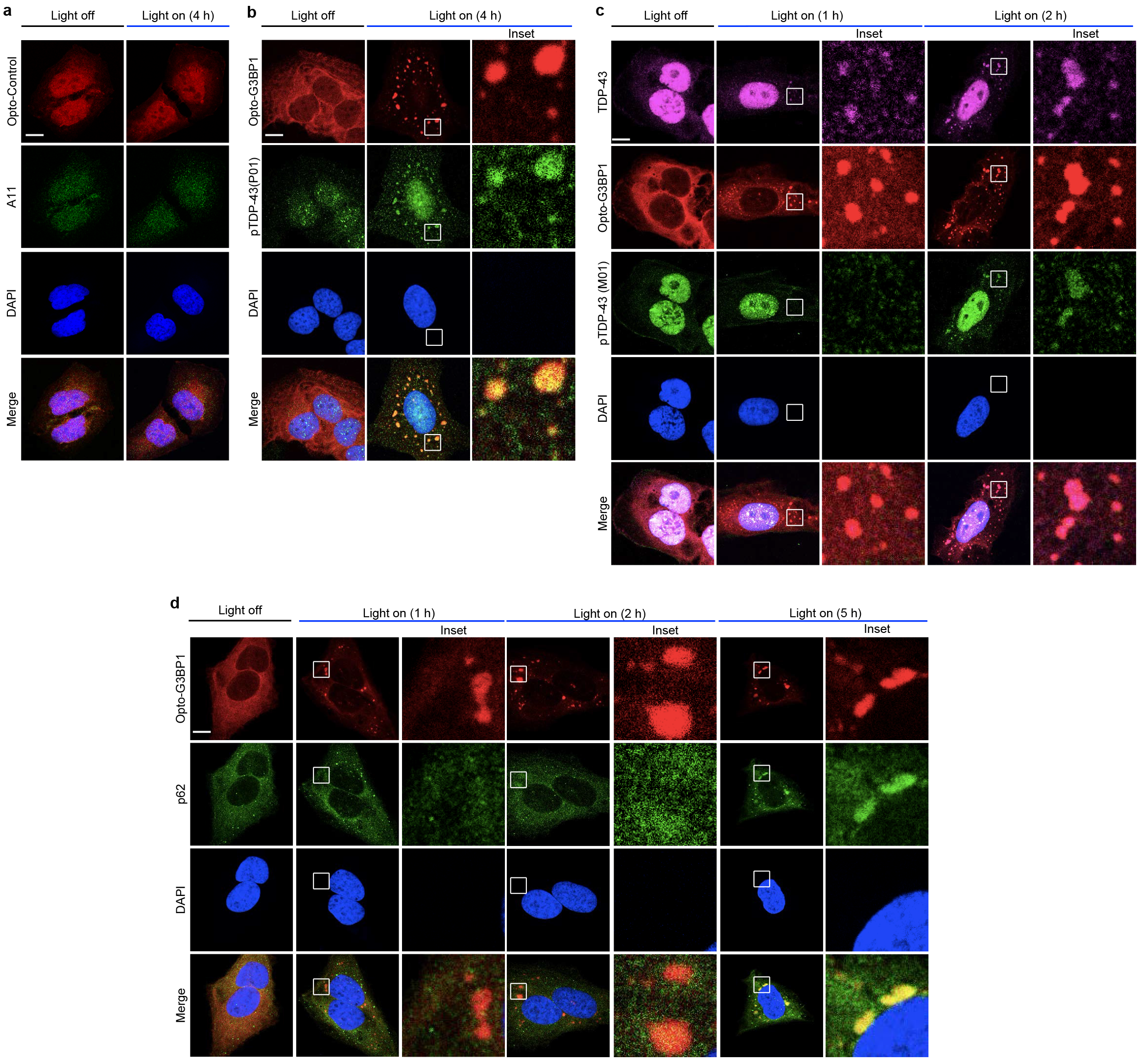
OptoGranules evolve to pathological inclusions. **a,** U2OS cells stably expressing Opto-Control were stimulated with blue light for 4 h and then immunostained for A11. Representative images are shown from n = 3 independent experiments. **b,** U2OS cells stably expressing Opto-G3BP1 were stimulated with blue light for 4 h and then immunostained for p-TDP-43 (p01). Representative images are shown from n = 2 independent experiments. **c-d,** U2OS cells stably expressing Opto-G3BP1 were stimulated with blue light for indicated times and then co-immunostained for TDP-43 and p-TDP-43 (M01) (**c**), or p62 (**d**). Representative images are shown from n = 3 independent experiments. Scale bars, 10 μm in all micrographs.

**Extended Data Figure 5.**
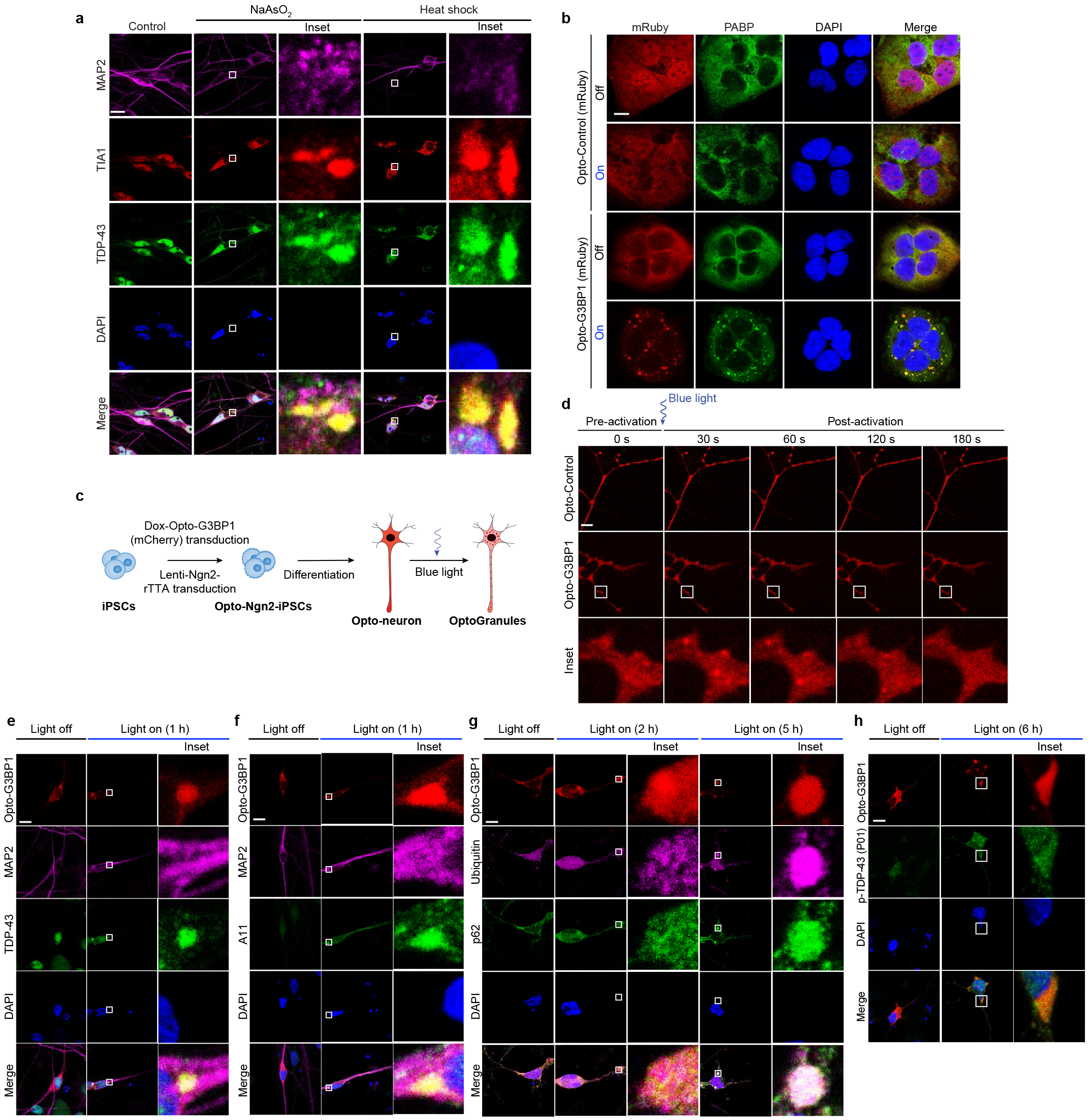
OptoGranules evolve to pathological inclusions in iPS cell-derived neurons. **a,** iPS cell-derived neurons expressing Opto-G3BP1 were treated with 0.25 mM sodium arsenite (NaAsO_2_) for 30 min or 42°C heat shock for 1 h and then immunostained for MAP2 along with stress granule marker TIA1 and TDP-43. **b,** U2OS cells stably expressing Opto-Control (mRuby) or Opto-G3BP1 (mRuby) were stimulated with blue light for 2 h and then immunostained for PABP. **c,** Schematic of doxycycline-inducible Opto-neuron generation. **d,** iPS cell-derived neurons expressing doxycycline-inducible Opto-Control (mCherry) or Opto-G3BP1 (mCherry) and stimulated with one 5-msec pulse of 488-nm blue light. Representative images are shown from n = 3 independent experiments. **e-h,** iPS cell-derived neurons expressing doxycycline-inducible Opto-G3BP1 were stimulated with blue light for indicated times and co-immunostained with MAP2 and TDP-43 antibodies (**e**), MAP2 and A11 antibodies (**f**), p62 and ubiquitin antibodies (**g**), or p-TDP-43 (P01) antibodies (**h**). Images in **e-h** are representative of n = 3 independent experiments. Scale bars, 10 μm in all micrographs.

## References

1 Kim, H. J. et al. Mutations in prion-like domains in hnRNPA2B1 and hnRNPA1 cause multisystem proteinopathy and ALS. Nature 495, 467–473, doi:10.1038/nature11922 (2013).

2 Lee, K. H. et al. C9orf72 Dipeptide Repeats Impair the Assembly, Dynamics, and Function of Membrane-Less Organelles. Cell 167, 774–788 e717, doi:10.1016/j.cell.2016.10.002 (2016).

3 Mackenzie, I. R. et al. TIA1 Mutations in Amyotrophic Lateral Sclerosis and Frontotemporal Dementia Promote Phase Separation and Alter Stress Granule Dynamics. Neuron 95, 808–816 e809, doi:10.1016/j.neuron.2017.07.025 (2017).

4 Molliex, A. et al. Phase separation by low complexity domains promotes stress granule assembly and drives pathological fibrillization. Cell 163, 123–133, doi:10.1016/j.cell.2015.09.015 (2015).

5 Patel, A. et al. A Liquid-to-Solid Phase Transition of the ALS Protein FUS Accelerated by Disease Mutation. Cell 162, 1066–1077, doi:10.1016/j.cell.2015.07.047 (2015).

6 Lin, Y. et al. Toxic PR Poly-Dipeptides Encoded by the C9orf72 Repeat Expansion Target LC Domain Polymers. Cell 167, 789–802 e712, doi:10.1016/j.cell.2016.10.003 (2016).

7 Lin, Y., Protter, D. S., Rosen, M. K. & Parker, R. Formation and Maturation of Phase-Separated Liquid Droplets by RNA-Binding Proteins. Mol. Cell 60, 208–219, doi:10.1016/j.molcel.2015.08.018 (2015).

8 Murakami, T. et al. ALS/FTD Mutation-Induced Phase Transition of FUS Liquid Droplets and Reversible Hydrogels into Irreversible Hydrogels Impairs RNP Granule Function. Neuron 88, 678–690, doi:10.1016/j.neuron.2015.10.030 (2015).

9 Buchan, J. R., Kolaitis, R. M., Taylor, J. P. & Parker, R. Eukaryotic stress granules are cleared by autophagy and Cdc48/VCP function. Cell 153, 1461–1474, doi:10.1016/j.cell.2013.05.037 (2013).

10 Becker, L. A. et al. Therapeutic reduction of ataxin-2 extends lifespan and reduces pathology in TDP-43 mice. Nature 544, 367–371, doi:10.1038/nature22038 (2017).

11 Boeynaems, S. et al. Phase Separation of C9orf72 Dipeptide Repeats Perturbs Stress Granule Dynamics. Mol. Cell 65, 1044–1055 e1045, doi:10.1016/j.molcel.2017.02.013 (2017).

12 Figley, M. D., Bieri, G., Kolaitis, R. M., Taylor, J. P. & Gitler, A. D. Profilin 1 associates with stress granules and ALS-linked mutations alter stress granule dynamics. J. Neurosci. 34, 8083–8097, doi:10.1523/JNEUROSCI.0543-14.2014 (2014).

13 Banani, S. F., Lee, H. O., Hyman, A. A. & Rosen, M. K. Biomolecular condensates: organizers of cellular biochemistry. Nat Rev Mol Cell Biol 18, 285–298, doi:10.1038/nrm.2017.7 (2017).

14 Kedersha, N. et al. G3BP-Caprin1-USP10 complexes mediate stress granule condensation and associate with 40S subunits. J. Cell Biol. 212, 845–860, doi:10.1083/jcb.201508028 (2016).

15 Kedersha, N. L., Gupta, M., Li, W., Miller, I. & Anderson, P. RNA-binding proteins TIA-1 and TIAR link the phosphorylation of eIF-2 alpha to the assembly of mammalian stress granules. J. Cell Biol. 147, 1431–1442 (1999).

16 Panas, M. D., Ivanov, P. & Anderson, P. Mechanistic insights into mammalian stress granule dynamics. J. Cell Biol. 215, 313–323, doi:10.1083/jcb.201609081 (2016).

17 Shin, Y. et al. Spatiotemporal Control of Intracellular Phase Transitions Using Light-Activated optoDroplets. Cell 168, 159–171 e114, doi:10.1016/j.cell.2016.11.054 (2017).

18 Sidrauski, C., McGeachy, A. M., Ingolia, N. T. & Walter, P. The small molecule ISRIB reverses the effects of eIF2alpha phosphorylation on translation and stress granule assembly. Elife 4, doi:10.7554/eLife.05033 (2015).

19 Neumann, M. et al. Ubiquitinated TDP-43 in frontotemporal lobar degeneration and amyotrophic lateral sclerosis. Science 314, 130–133, doi:10.1126/science.1134108 (2006).

20 Mackenzie, I. R. et al. Pathological TDP-43 distinguishes sporadic amyotrophic lateral sclerosis from amyotrophic lateral sclerosis with SOD1 mutations. Ann Neurol 61, 427–434, doi:10.1002/ana.21147 (2007).

21 Mackenzie, I. R. & Neumann, M. Molecular neuropathology of frontotemporal dementia: insights into disease mechanisms from postmortem studies. J Neurochem 138 Suppl 1, 54–70, doi:10.1111/jnc.13588 (2016).

22 Williams, K. L. et al. UBQLN2/ubiquilin 2 mutation and pathology in familial amyotrophic lateral sclerosis. Neurobiol Aging 33, 2527 e2523–2510, doi:10.1016/j.neurobiolaging.2012.05.008 (2012).

23 Deng, H. X. et al. Mutations in UBQLN2 cause dominant X-linked juvenile and adult-onset ALS and ALS/dementia. Nature 477, 211–215, doi:10.1038/nature10353 (2011).

24 Neumann, M. et al. Phosphorylation of S409/410 of TDP-43 is a consistent feature in all sporadic and familial forms of TDP-43 proteinopathies. Acta Neuropathol 117, 137–149, doi:10.1007/s00401-008-0477-9 (2009).

25 Zhang, K. et al. Stress Granule Assembly Disrupts Nucleocytoplasmic Transport. Cell 173, 958–971 e917, doi:10.1016/j.cell.2018.03.025 (2018).

26 Zhang, Y. et al. Rapid single-step induction of functional neurons from human pluripotent stem cells. Neuron 78, 785–798, doi:10.1016/j.neuron.2013.05.029 (2013).

27 Richner, M., Victor, M. B., Liu, Y., Abernathy, D. & Yoo, A. S. MicroRNA-based conversion of human fibroblasts into striatal medium spiny neurons. Nat. Protoc. 10, 1543–1555, doi:10.1038/nprot.2015.102 (2015).

## References for Methods

1 Zhang, K. et al. Stress Granule Assembly Disrupts Nucleocytoplasmic Transport. Cell 173, 958–971 e917, doi:10.1016/j.cell.2018.03.025 (2018).

2 Figley, M. D., Bieri, G., Kolaitis, R. M., Taylor, J. P. & Gitler, A. D. Profilin 1 associates with stress granules and ALS-linked mutations alter stress granule dynamics. J. Neurosci. 34, 8083–8097, doi:10.1523/JNEUROSCI.0543-14.2014 (2014).

4 Lee, K. H. et al. C9orf72 Dipeptide Repeats Impair the Assembly, Dynamics, and Function of Membrane-Less Organelles. Cell 167, 774–788 e717, doi:10.1016/j.cell.2016.10.002 (2016).

5 Zhang, Y. et al. Rapid single-step induction of functional neurons from human pluripotent stem cells. Neuron 78, 785–798, doi:10.1016/j.neuron.2013.05.029 (2013).

6 Richner, M., Victor, M. B., Liu, Y., Abernathy, D. & Yoo, A. S. MicroRNA-based conversion of human fibroblasts into striatal medium spiny neurons. Nat. Protoc. 10, 1543–1555, doi:10.1038/nprot.2015.102 (2015).

7 Valentin-Vega, Y. A. et al. Cancer-associated DDX3X mutations drive stress granule assembly and impair global translation. Sci. Rep. 6, 25996, doi:10.1038/srep25996 (2016).

